# biGMamAct: efficient CRISPR/Cas9-mediated docking of large functional DNA cargoes at the *ACTB* locus

**DOI:** 10.1101/2024.04.18.590029

**Authors:** Martin Pelosse, Marco Marcia

## Abstract

Recent advances in molecular and cell biology and imaging have unprecedentedly enabled multi-scale structure-functional studies of entire metabolic pathways from atomic to micrometer resolution, and the visualization of macromolecular complexes *in situ*, especially if these molecules are expressed with appropriately-engineered and easily-detectable tags. However, genome editing in eukaryotic cells is challenging when generating stable cell lines loaded with large DNA cargoes. To address this limitation, here, we have conceived biGMamAct, a system that allows the straightforward assembly of a multitude of genetic modules and their subsequent integration in the genome at the *ACTB* locus with high efficacy, through standardized cloning steps. Our technology encompasses a set of modular plasmids for mammalian expression, which can be efficiently docked into the genome in tandem with a validated Cas9/sgRNA pair through homologous-independent targeted insertion (HITI). As a proof of concept, we have generated a stable cell line loaded with an 18.3-kilobase-long DNA cargo to express 6 fluorescently-tagged proteins and simultaneously visualize 5 different subcellular compartments. Our protocol leads from the *in-silico* design to the genetic and functional characterization of single clones within 6 weeks and can be implemented by any researcher with familiarity with molecular biology and access to mammalian cell culturing infrastructure.

## Introduction

Integration of structural and cellular biology has recently enabled the characterization of large and complex systems at different resolution scales, *in situ*^1,2^. Studying biological macromolecules within their cellular environment enables the visualization of vital biological processes in physiological conditions, and consequently improves our understanding fundamental biological mechanisms. However, *in situ* molecular characterization faces specific challenges and requires new technological development.

An important challenge when studying multi component macromolecular systems is the number of genetic modules that need to be engineered, assembled, and expressed in the cell. For generating stable cell lines, assembling genetic cassettes in a way compatible with genomic integration is a limiting factor. Available cloning systems designed for easy and rapid assembly of multiple DNA elements are meant to be used insect cells and can rarely be deployed in mammalian cells^3,4^. Moreover, despite their robustness, specific skills in molecular biology, such as site-specific recombineering or multisite gateway cloning, are required to establish such tools^3,5–7^, which limits their application.

Precise gene editing has become possible thanks to techniques such as CRISPR/Cas^8^, TALEN^9,10^, or Prime Editing^11^, which allow to alter endogenous genes (KO, targeted mutation, tagging, cure of genetic disease^12,13^). Generation of stably-edited pools and clonal subpopulation enables researchers to work with homogenous cell samples and can unlock the study of subtle biological process, even at the single cell level. However, pitfalls and limitation are still profuse when the question at issue applies to the experimental design using these nucleases. Indeed, TALEN entails a complicated proteins design^14,15^ and validation. CRISPR/Cas9 is limited by sgRNA selection and validation^16,17^ or template design inherent to the choice of the repair mechanism exploitable in given cell types (dividing vs. non-dividing). Furthermore, when working in living systems, inherent variability is increased by some other time-related limitations that should also be considered, such as silencing^18^.

To address these limitations and maximize the potential of currently-available genetic engineering technology, here, we have designed a method for multi components assembly and tunable expression in mammalian cells, with highly efficient and precise CRISPR/Cas9 mediated insertion into the genome. Our method will pave the way for easily and quickly generating stable cells lines and facilitate the study of complex biological processes at a multiscale cellular level, especially those requiring the simultaneous visualization of multiple macromolecules^19^.

### Development of the method

In our facility, we extensively use MultiBac^20^ and biGBac^21^ baculovirus expression vector systems (BEVS) for assembling large DNA constructs encoding for large multiprotein complexes, their subsequent expression in insect cells and structure determination^22,23^. Because biGBac uses standardized bioinformatically-validated DNA sequences for ligation independent cloning (LIC)^24,25^ and minimizes the carry-over of noncoding sequences (plasmid backbone) during sequential assemblies, we decided to adapt it for deployment in mammalian cells.

When tackling delivery issues inherent to increasing size of DNA payloads for genomic integration in specific cell lines (by involving engineered baculovirus^26^), we validated a highly efficient way of knocking in foreign DNA at the actin-β (*ACTB*)^4^ locus through homology-independent targeted insertion (HITI)^27^ consecutive to genomic Cas9-induced double strand break.

Here, we fused these two methodologies to design a robust tool for easy and quick assembly of several genetic module into a large construct, which can be efficiently and precisely integrated in both dividing and non-dividing cells. Our method relies on established molecular biology skills accessible to any researcher, and builds on straightforward experimental design, relying on our already established *ACTB* targeting by CRISPR/Cas9.

Using our novel methodology scientists have the possibility to generate stable cell lines with precisely-inserted functional DNA cargo of several tens of kilobases, expressing tens of macromolecular subunits (here, we benchmarked the protocol by inserting an 18.3 kb-long DNA cargo expressing 6 fluorescent proteins to simultaneously visualize 5 different cellular compartments).

### Overview of the procedure

Our new method offers the possibility to i) easily assemble in one standardized step, through LIC, up to 6 genetic modules which in turn can be combined with the use of self-cleaving peptides^28^ and ii) to reliably and stably integrate these modules into the genome at the *ACTB* housekeeping locus using a CRISPR/Cas9-based approach^4^. Importantly, our assembly system preserves the highly modular properties of the previously-reported biGBac system, hence up to 25 modules can be assembled in only 2 standardized LIC steps to create a single all-in-one carrier vector^21^.

To overcome working with several plasmids, often leading to biased results due to variable DNA uptake by the cells^29^, it is recommended to generate one large single DNA vector encompassing all genes of interest. Because this approach can sometimes involve tedious cloning methodologies, we decided to adapt the reliable and standardized biGBac cloning system towards mammalian cells, thus generating our new biGMamAct system. Through our approach, within a week any scientist can assemble in one single step up to 6 genetic modules leading to the generation of an all-in-one coding plasmids for subsequent integration at the *ACTB* locus (**Fig 1** and **Fig 2A**). To do so, assembled plasmid (pDONOR), will be co-transfected with standard pX458*ACTB*, encoding for spCas9 and sgRNA targeting last intron of the human *ACTB* locus. Thanks to our design, spCas9 will not only induce a double strand break in the genome but will also linearize the pDONOR allowing its stable and precise integration at the *ACTB* locus (**Fig 2A**). After 2 weeks of double selection applied onto the transfected cells, the desired stable cell line is established and 2 more weeks are required for selection and genomic validation of clonal subpopulations (**Fig 1**).

**Figure 1:**
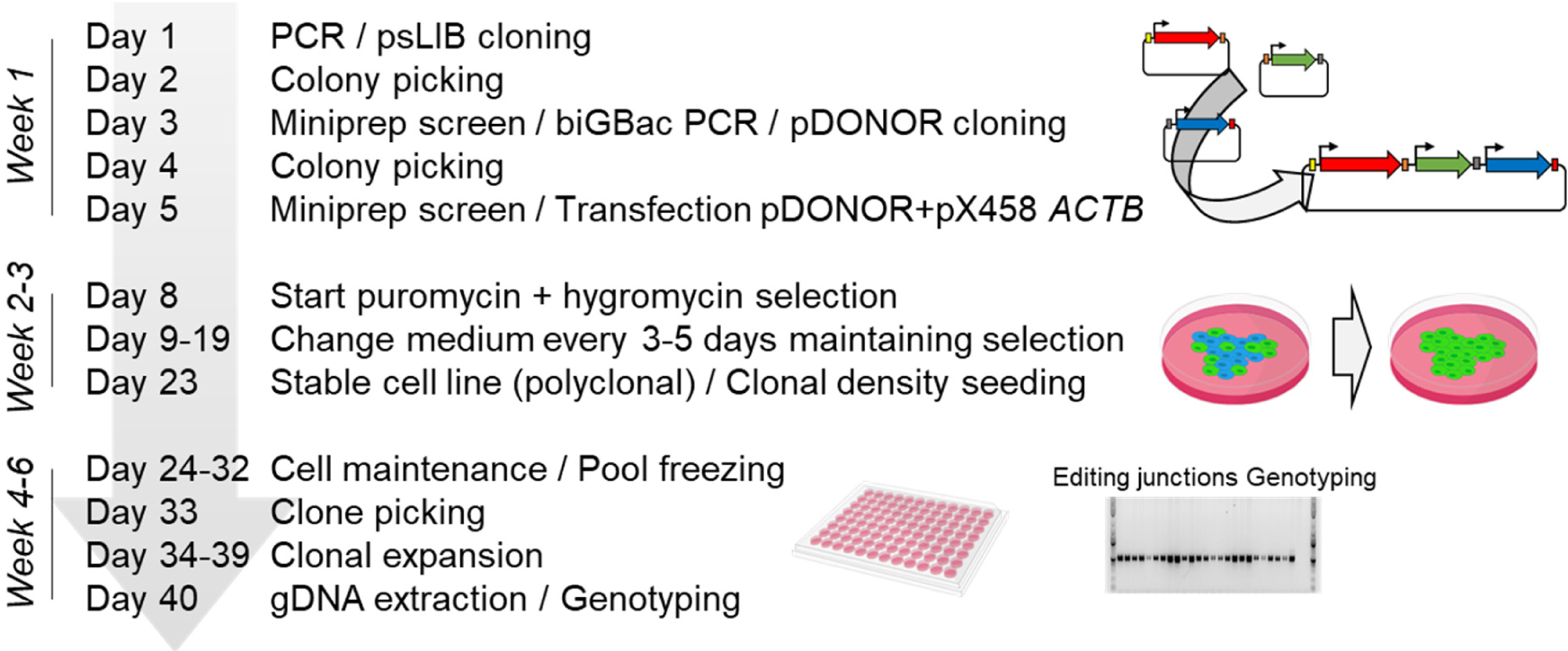
Workflow of the biGMamAct method. The first week of cloning is followed by 2 weeks of selection to establish a polyclonal stable cell line. 2 more weeks are required to expend, isolate and genotype clonal population *(partially designed using bioicons.com)*.

**Figure 2:**
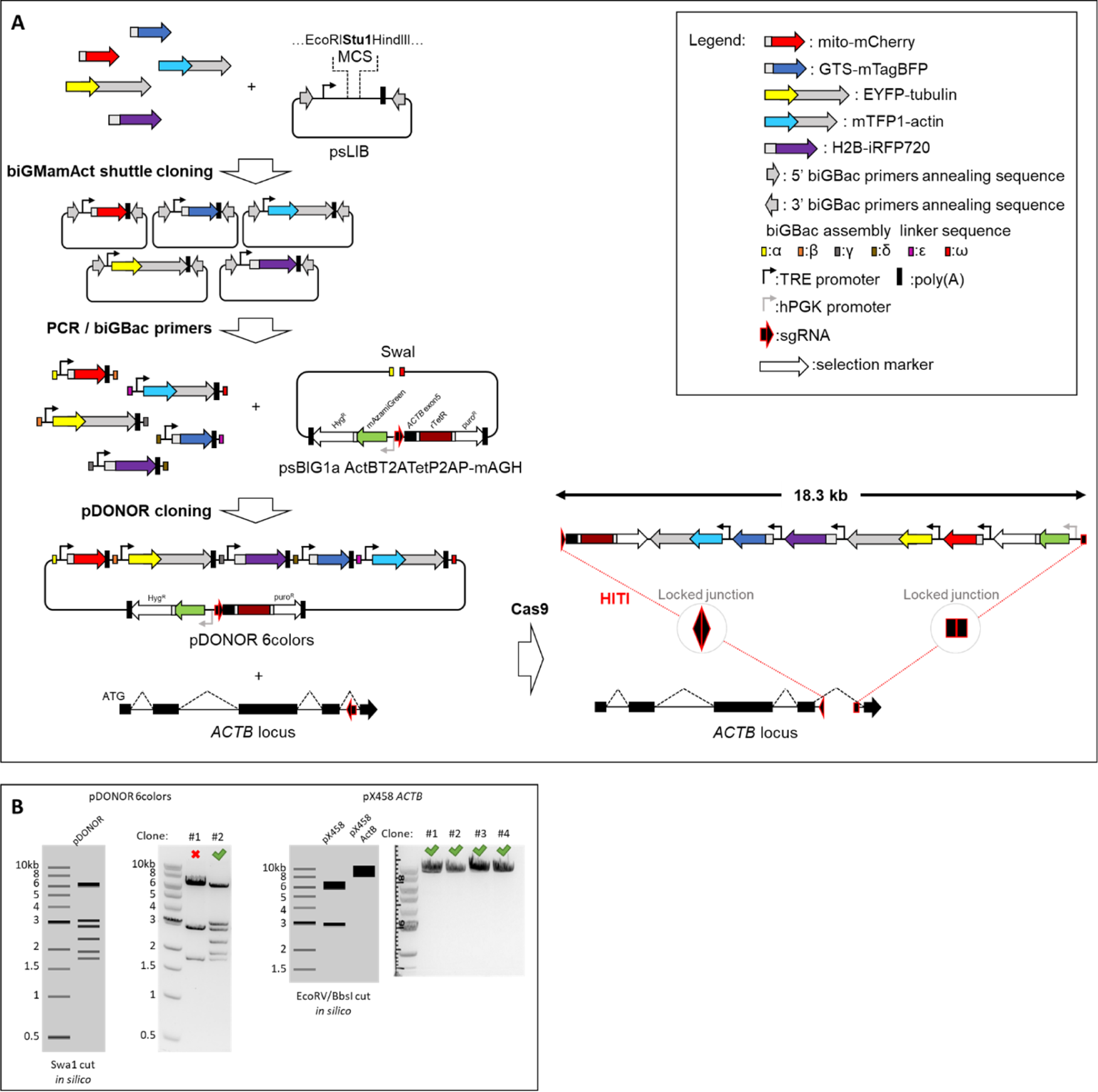
A) biGMamAct assembly and knockin design. Genes of interest (GOI) inserted into the psLIB shuttle plasmids of the biGMamAct expression system can be PCR amplified using the standard bioinformatically designed primers of the biGBac system for further assembly into our recipient psBIG1a *ACTB*T2ATetP2AP-mAGH. Co-transfection of the pDONOR 6colors plasmid with a Cas9-containing-one targeting *ACTB* locus will lead to pDONOR stable and precise integration via HITI at the *ACTB* locus with high efficiency. Indeed, *ACTB* sgRNA targeted sequence is present in reverse orientation in pDONOR for linearization by Cas9 endonuclease and orientation-controlled integration in the genome. **B) Cloning validation.** Agarose gels showing recombining and positive pDONOR 6colors clones (left) and correct sgRNA *ACTB* cloning in pX458 (right).

### Application of the method

We validated our method by generating stable cell lines harboring inducible tunable reporters allowing simultaneous tagging of 5 different subcellular compartments, i.e. mitochondria, the Golgi, the cytoskeleton (actin), microtubules (tubulin), and the nucleus. We envisage that our method can be used to generate cell lines, as efficient cell factory, stably loaded with the components of entire metabolic pathways^30^, which can be turned on and off through the use of our tunable promoter (**Table 1**). BiGMamAct will certainly quickly integrates the synthetic biology tools for engineering mammalian cell systems and match some required needs in this emerging field^31–33^. Our methods will certainly also open new avenues for *in situ* structural biology studies providing the opportunity to integrate into cells tools for localization of macromolecular complexes^34,35^.

**Table 1:**
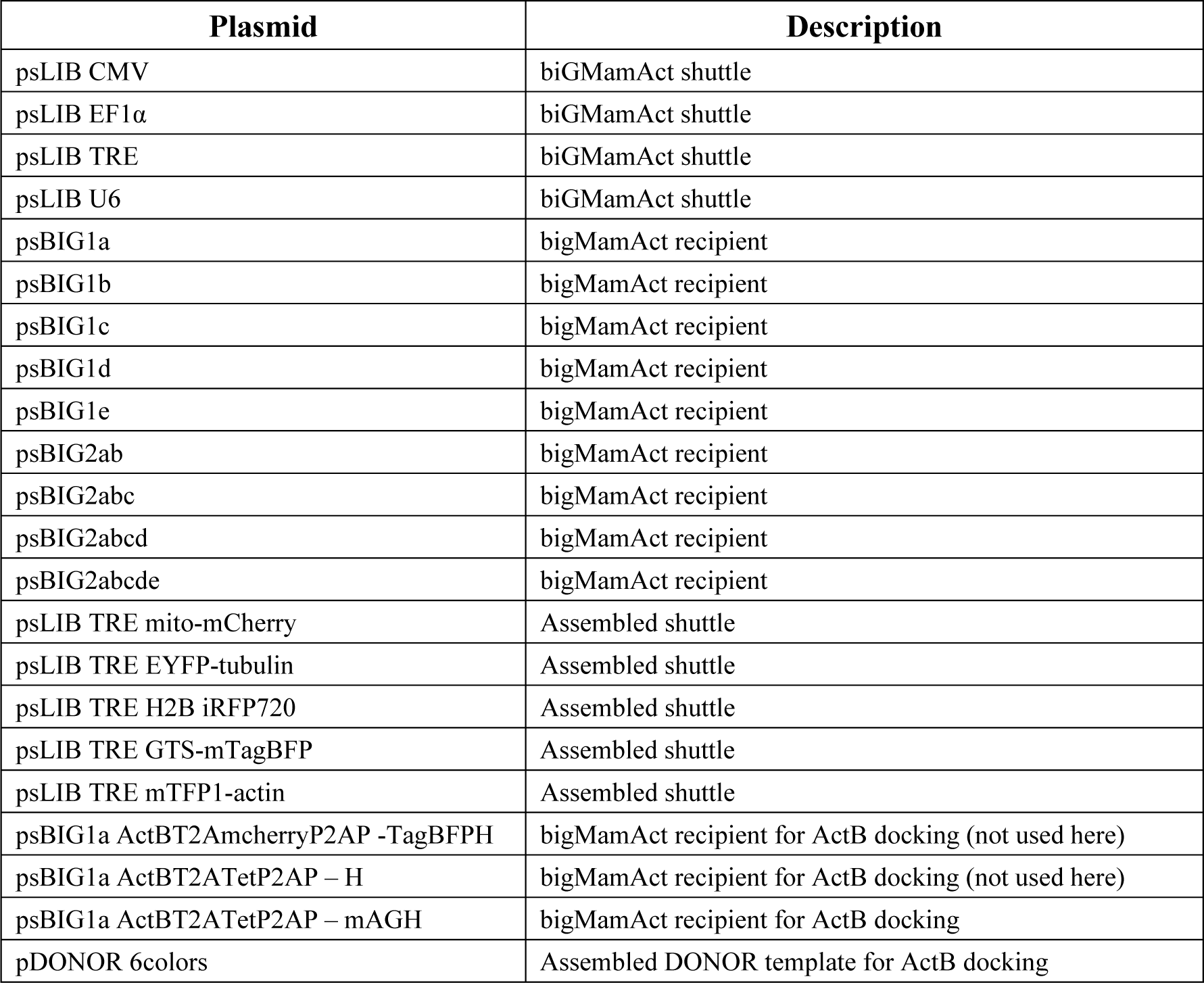
biGMamAct Plasmid library.

### Comparison with other methods /advantages

Our method exploits the use of biGBac standardized and bioinformatically-validated oligos for cloning and a well-established genome engineering design for quick and efficient generation of stable cell lines harboring large functional DNA payloads. Indeed, in our hands biGBac is the cloning system allowing the easiest assembly of a large number of genetic modules. Moreover, genomic integration though HITI after a Cas9 double strand break is the most efficient reported approach for knocking in foreign DNA into the genome of any cell types^4,36^.

Using a single all-in-one plasmid offers the chance to overcome uncontrolled stoichiometry of cellular uptake of plasmids, which would lead to artefactual results or experimental failure. Some cloning systems have been designed for easy and rapid assembly of multiple DNA elements and are available. However, most of them are part of Baculovirus Expression Vector System (BEVS) and solely intended to be used for multiprotein complexes expression in insect cells^20,21,37^. Very few can be deployed in mammalian cells^3,4^ and despite their robustness and efficacy they remain isolated in use as requiring extensive expertise in molecular biology. Multigene assembly using MultiMAM is achieved via Cre/Lox recombination and MuliMATE requires another recombinatorial cloning, multisite Gateway. Here, instead, as part of our new biGMamAct technology, we have designed and generated an optimized version of the biGBac cloning system suitable for expression in mammalian cells and compatible with CRISPR/Cas mediated insertion at the *ACTB* locus.

Leaning on CRISPR/Cas9 precise integration instead of random integration enables the integration of donor sequence as a whole. Indeed, if large DNA payload are considered for genomic introduction, random integration could instead be limited to surrounding sequences of the selection markers. CRISPR/Cas9 mediated docking of all-in-one pDONOR also reduces the number of selection markers required when using several plasmids. Our method relies on standardized oligos for cloning and a validated spCas9/sgRNA system targeting *ACTB* to ensure high efficiency and time saving^4,38^. Indeed, other approaches targeting other loci would require sgRNA design and validation. Instead, other approaches involving nucleases such as TALEN would necessitate tedious, cloning and validation before being applicable.

Furthermore, other available methods can be used for smaller DNA precise integration such as PrimeEditing^39^ but are not suitable for large DNA cargo. PiggyBac transposon-based methods^40^ or lentivirus can also be used for stable integration in genomes. However, these are lacking precision regarding their targeted sequences and cannot be adapted to DNA payloads considered herein. Most of the current methods for precise genome editing rely on Homology Direct Repair (HDR) following a CRISPR/Cas induced double strand break. It has to be acknowledged that repair through Non-Homologous End Joining (NHEJ) is leading to integration of small insertion deletions (indels) at the cut site. Genome editing for therapeutic purpose or in patients should thus avoid being based on such DNA repair pathway. However, it is known that NHEJ is occurring in dividing and non-dividing cells, contrary to HDR, and methods based on NHEJ, such as HITI were reported to show greater absolute efficiency of 1 to 2 order of magnitude than HDR-based ones, and applicable in any cell type. Relying on both CRISPR/Cas9 and HITI as cut- and insertion mechanisms enable us to precisely integrate up to 18.2 kb functional DNA payload with 100% of the screened clones being homozygous (24/24) (**Fig 3**).

**Figure 3:**
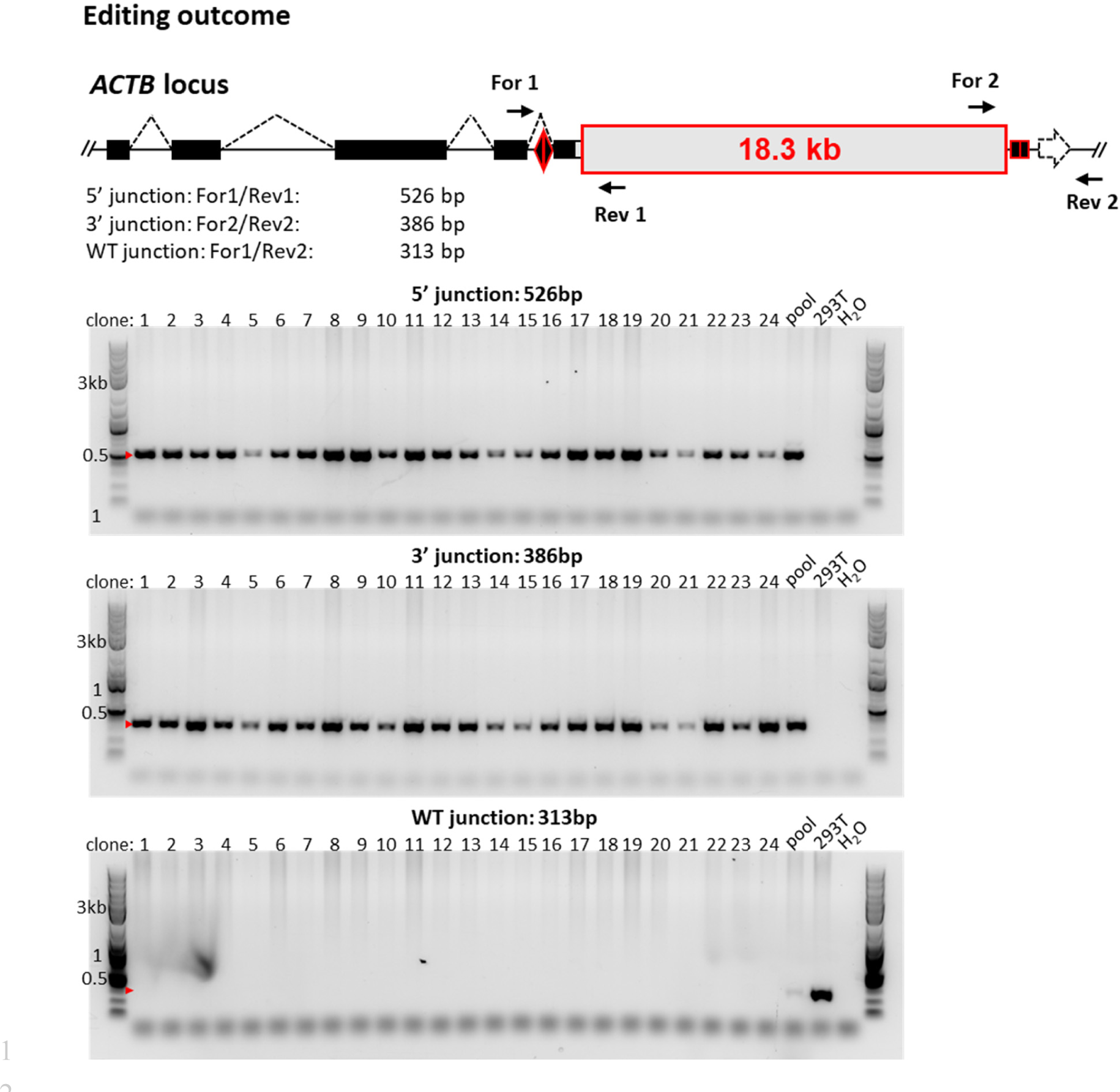
Genotyping of clone highlight biGMamAct high efficiency knock-in at *ACTB* loci. Schematic representation of edited human *ACTB* locus. Black triangle and boxes encircled in red represents the locked 5’ and 3’junction resulting from HITI docking. Primers used for genotyping are depicted as black arrows surrounding these junctions. Correct DNA cargo integration lead to visualization of bands at 526 bp and 386 bp for the 5’ and 3’ junction respectively. Heterozygotes still show a band at 313bp outlining the persistence of the WT junction (only in pool and HEK293T WT).

Finally, silencing over time is another important limitation that biologists are facing when generating stable cells. Here we combine targeting of the *ACTB* locus, a constantly processed gene over the whole cell cycle, to inducible and tunable promoters to circumvent this issue and offering as another advantage, to be able to work with toxic proteins.

In summary, with respect to available genetic engineering technologies, our method unprecedentedly overcomes three important limitations, such as the difficulty in generating large DNA cargoes, the precision of integration, and the constitutive expression of the genes of interests.

### Limitations of the method

We can identify four possible limitations in our method.

First, NHEJ introduces small indels which, in our case, are confined to the last intron of the *ACTB* gene. It has to also be mentioned that our method is not replacing the last *ACTB* exon by a fusion one but simply shifting it to the 3’UTR, representing another mark induced by our editing method.

Second, the establishment of a stable cell line following our method uses 2 selection markers and lasts 15 days, which can be long when working with primary cells. To overcome this limitation, we recommend swapping selection markers for fluorescent ones combined to fluorescence activated cell sorting (FACS).

Third, our procedure relies on lipofection of the pDONOR and Cas9+sgRNA containing plasmids. This might not be ideal considering given cell types for which nucleofection should be favored. However, our system is fully compatible with other transfection methods and can still be deployed in cells showing low transfection efficiency as HITI shows the highest precise integration efficiency.

Fourth, our method carries one limitation inherent to the high modularity of the biGBac system^21^. Indeed, assembling dozens of genetic modules rises the problem of recombineering during the first steps of cloning. To avoid plasmid recombination due presence of repetitive elements, we recommend taking advantage of self-cleaving peptides during the initial construct design and propose some tips for limiting such issue. In the present study, the all-in-one vector assembled contains 40 repeats of the Tet operator sequence suggesting recombination related issues are not a major limitation of the method, if at all.

### Rational design

#### Plasmids engineering

Working with several isolated vectors may lead to uncontrolled variability when considering relative transfection efficiency of individual plasmids and stochiometric recombinant expression of encoded proteins inside cells. The biGMamAct system encompasses a set of plasmids derived from the biGBac cloning system^21^. In the present study, we considerably reduced in size (32% reduction) the existing pLIB and created a smaller version, psLIB. The psLIB backbone still allows single cassette amplification by PCR using the standardized and bioinformatically designed set of biGBac primers. The polH promoter for insect cells was replaced by various promoters to match user needs (CMV, EF1α or Tet responsive element (TRE)) (**Table 1**). The possibility to use TRE as promoter allows to work with cytotoxic proteins and also to limit silencing possibly occurring while establishing the stable cell line. Up to 5 PCR-amplified cassettes from psLIB can be inserted at once by Gibson cloning^25^ (LIC) into our psBIG1a *ACTB*T2ATetP2AP-mAGH harboring the biGBac α and ω sequences. The psBIG1a *ACTB*T2ATetP2AP-mAGH also carries biGBac A and B sequences, thus cassettes cloned into it can further be combined with others inserted into psBIG1 plasmids (b to e) through another step of standardized Gibson cloning into psBIG2 plasmids (ab to abcde). psBIG1 and psBIG2 plasmids series are 28% smaller than the pBIG1 and pBIG2 from biGBac^21^ allowing easier manipulation and/or increasing loading capacity. Similar to biGBac, these are up to 25 independent cassettes which can be assembled in only two cloning steps as exemplified in the original study^21^. However, we recommend also making use of self-cleaving peptides as a way of reducing the number of individual cassettes. This strategy will limit the number of repeated sequences (e.g.: promoters) which could lead to unwanted recombination events during the cloning. Importantly, as part of the biGMamAct cloning system, the newly created psLIB_CMV and psLIB_EF1α can be used and assembled together with the smaller psBIG1 and psBIG2 vectors for transient expression of dozens of independent cassettes. biGMamAct plasmids psBIG1 and psBIG2 offer the possibility to insert generated very large construct into baculovirions for high transduction efficiency in a wide variety of cell types^4,26,38,41^.

The psBIG1a *ACTB*T2ATetP2AP-mAGH contains the same sgRNA sequence than the *ACTB* locus targeted (**Fig 2A**). This feature implies that we exploit Cas9 for linearization of the exogenous DNA to be knocked-in (pDONOR). The psBIG1a *ACTB*T2ATetP2AP-mAGH sequence also have two selection markers (puromycin and hygromycin) surrounding the sgRNA sequence to produce, once being cut, the DNA cargo of interest being flanked by two selection markers. Through this design our method ensures complete integration of the DNA cargo at the correct genomic location. Being able to apply selection pressures at both 5’ and 3’-ends of the cargo DNA further limits the chance of silencing by keeping the genomic region active. If cell sorters are available, fluorescent proteins can easily be inserted or removed as fusion constructs with the selection markers genes. The use of double selection markers, which can be combined to the use of two fluorescent markers (e.g. mCherry and mTagBFP-NLS or mAzamiGreen(mAG)-NLS), allows for efficient and reliable selection during stable cell line generation and clonal characterization.

#### Choice of integration mechanism

When a double strand cut occur into the genome, an exogenous piece of DNA can be introduced through two independent mechanisms: Homology Targeted Repair (HDR) or Non-Homologous End Joining (NHEJ), each one relying on a different set of proteins^42,43^. For making our method the more universal and as efficient as possible we decided to rely on NHEJ-repair mechanisms and to involve the powerful and underrated Homology-Independent Targeted Insertion (HITI)^27^. Indeed, HITI has been reported to be the most efficient strategy to mediate knock-ins^27,44^. In addition, and in contrast to low efficiency HDR, HITI was shown to occur at every stage of cellular life, rendering our method applicable into dividing and non-dividing cells. This also has for advantage to abolish the needs of long homology arms to be cloned into the donor plasmids increasing the adaptability of our method in the case of targeting of other genomic loci (sgRNA sequence exchange only required) and increasing both the cargo capacity and the transfection efficiency of pDONOR. By introducing into our pDONOR plasmid, the same targeted sequence (sgRNA+PAM) than the one by spCas9 in the genome, but in reverse orientation, we exploit spCas9 to linearize the DNA to be inserted, whilst ensuring its correct integration and orientation (**Fig 2A**). Indeed, pDONOR insertion in reverse orientation would result in restauration of the sgRNA+PAM sequence leading to spCas9 cleavage again and excision of the wrongly inserted pDONOR (**Fig 2A**). Because HITI is not a scar-free integration mechanism we have picked the best sgRNA within the last intron of *ACTB* by combining use of E-CRISP^45^ and IDT’s gRNA design checker. Again, making use of the NHEJ-dependent mechanism, HITI, subsequent to spCas9 cuts into both the genome and pDONOR underlined the power of the method described here, as we demonstrate precise integration of functional DNA of unprecedented size with 100 % of the screened clones being homozygotes (**Fig 3**).

#### Choice of locus and selection

In our method, we have selected the *ACTB* locus as landing pad for our large DNA cargo. Actin β is an essential gene being highly expressed all along the cell cycle. Moreover, this locus has been reported to be a very good candidate for docking exogenous DNA sequence for stable cell line generation. Indeed, it has been reported to be less - or equally - prone to silencing than other well-established docking sites^46,47^. To further reduce the risk of silencing over time, we integrated into our portfolio of shuttle plasmids some harboring Tet-inducible promoters. Picking *ACTB* as targeted genomic locus also makes the present technology applicable into already established cells lines relying on AAVS1^48^, hRosa26^49^, CCR5 and others^50^. However, out method can easily be further deployed at other genomic loci only by using other selection markers and sgRNAs (housekeeping loci e.g. GAPDH or safe harbor sites e.g. AAVS1, or others^50,51^).

Our method targets the last *ACTB* intron and, when inserting the foreign DNA, restores the final exon as a fusion construct harboring an in-frame puromycin selection marker. By this approach, our method abolishes off-targets insertion that could be observed depending on sgRNA sequences and when using highly-processive spCas9. Thanks to the 2 selection markers flanking the *ACTB* sgRNA+PAM sequence in pDONOR, the cleavage by spCas9 nuclease produces a 5’-end fusion of the last *ACTB* exon with puromycin resistant marker and a 3’-end consisting in a hygromycin selection cassette. Applying both selections will thus ensure precise and complete integration of the whole pDONOR cargo.

## Materials

### Reagents

**! CAUTION** We advise wearing protective clothing (lab coats and gloves) when handling any reagent mentioned in this protocol. Recycling or disposal of solid and liquid waste should be done according to local and institutional regulations

**▴CRITICAL** All reagents should be stored and prepared according to the manufacturer’s recommendations, if not otherwise indicated.

- Luria Bertani (LB) medium (Dutscher, cat. no. 777495)
- Agar (Euromedex France, cat. no. 1329-D)
- Agarose (Euromedex, cat. no. D5-D)
- TAE buffer 1X (40 mM Tris, 20 mM acetic acid, 1mM EDTA):

- Tris base (Euromedex France, cat. no. 200923-A) **! CAUTION** This reagent irritates eyes, respiratory system, and skin. Avoid all contact with eyes and skin and handle the reagent under a laminar flow hood to avoid inhalation.
- EDTA (Carl Roth, cat. no. 80431) **! CAUTION** This reagent irritates the eyes. Avoid all contact with eyes.
- Acetic acid (VWR, cat. no. 20104.298) **! CAUTION** This reagent is flammable and causes severe burns. Handle the reagent away from flames
- Top10 chemically competent cells (ThermoFisher, cat. no. C404003) **▴CRITICAL** Store the cells at −80 °C for up to 1 year.
- HEK293T cells (ATCC CRL 3216)
- Gentamycin (ThermoFisher, cat. no. J62834.06) **! CAUTION** This reagent irritates eyes, respiratory system, and skin and may cause sensitization upon inhalation and skin contact. Avoid all contact with eyes and skin, and handle the reagent under a laminar flow hood to avoid inhalation
- Puromycin (ThermoFisher, cat. no. A1113803)
- Hygromycin (ThermoFisher, cat. no. 10687010) **! CAUTION** Very toxic and carcinogenic substance that may cause death or damage to health when inhaled, swallowed or absorbed through the skin in small quantities. Wear protective gloves/protective clothing/eye protection/face protection.
- DpnI enzyme (New England Biolabs, cat. no. R0176S)
- StuI enzyme (New England Biolabs, cat. no. R0187S)
- SwaI enzyme and Neb3.1 buffer (New England Biolabs, cat. no. R0604S)
- biGBac set of primers (Table 2 and ^21^)
- Q5® Hot Start High-Fidelity DNA Polymerase (New England Biolabs, cat. no. M0493)
- KAPA2G Fast HotStart ReadyMix (Roche, cat no. 2GFHSGKB)
- dNTPs (New England Biolabs, cat. no. N0447S)
- Milli-Q water (ultrapure water, 18.2 MΩꞏcm at 25 °C)
- NEBuilder® HiFi DNA Assembly Master Mix (New England Biolabs, cat. no. E2621S)
- Gel Loading Dye, Orange (6X) (New England Biolabs, cat. no. B7022S)
- 1kb Plus DNA ladder (New England Biolabs, cat. no. N3200S)
- GelRed® (Merck, cat. no. SCT123)
- Scalpel blade and tweezer for DNA band excision and removal **! CAUTION** Make sure to work slowly and carefully
- QIAprep Spin Miniprep Kit (Qiagen, cat. no. 27106)
- QIAquick Gel Extraction Kit (Qiagen, cat. no. 28704)
- Monarch® PCR & DNA Cleanup Kit (New England Biolabs, cat. no. T1030S)
- Monarch® Genomic DNA Purification Kit (New England Biolabs, cat. no. T3010S)
- Dulbecco’s Modified Eagle Medium (DMEM) (ThermoFisher, cat. no. 41965062)
- Fetal Bovine Serum (FBS) (ThermoFisher, cat. no. 26140079)
- Penicillin/streptomycin (10 000 U/ml) (ThermoFisher, cat. no. 15140122) **! CAUTION** Suspected of damaging fertility or the unborn child. Wear protective gloves/protective clothing/eye protection/face protection.
- Tryspsin-EDTA (0.25%) (ThermoFisher, cat. no. 25200056)
- LipoD293 (Signagen, cat. no. SL100668)
- PBS (ThermoFisher, cat. no. 14190250)
- Dimethyl Sulfoxide (DMSO) sterile (Merck, cat. no. D2650)
- PFA 4% (ThermoFisher, cat. no. J19943.K2) **! CAUTION** Very toxic and carcinogenic substance that may cause death or damage to health when inhaled, swallowed or absorbed through the skin in small quantities. Wear protective gloves/protective clothing/eye protection/face protection.
- ProLong™ Diamond Antifade Mountant with DAPI (ThermoFisher, cat. no. P36966)
- Ethanol (VWR, cat. no. 20816.367) **! CAUTION** This reagent is highly flammable. Handle the reagent away from flames.
- Collagen (Roche, cat. no. 11179179001)

**Table 2:**
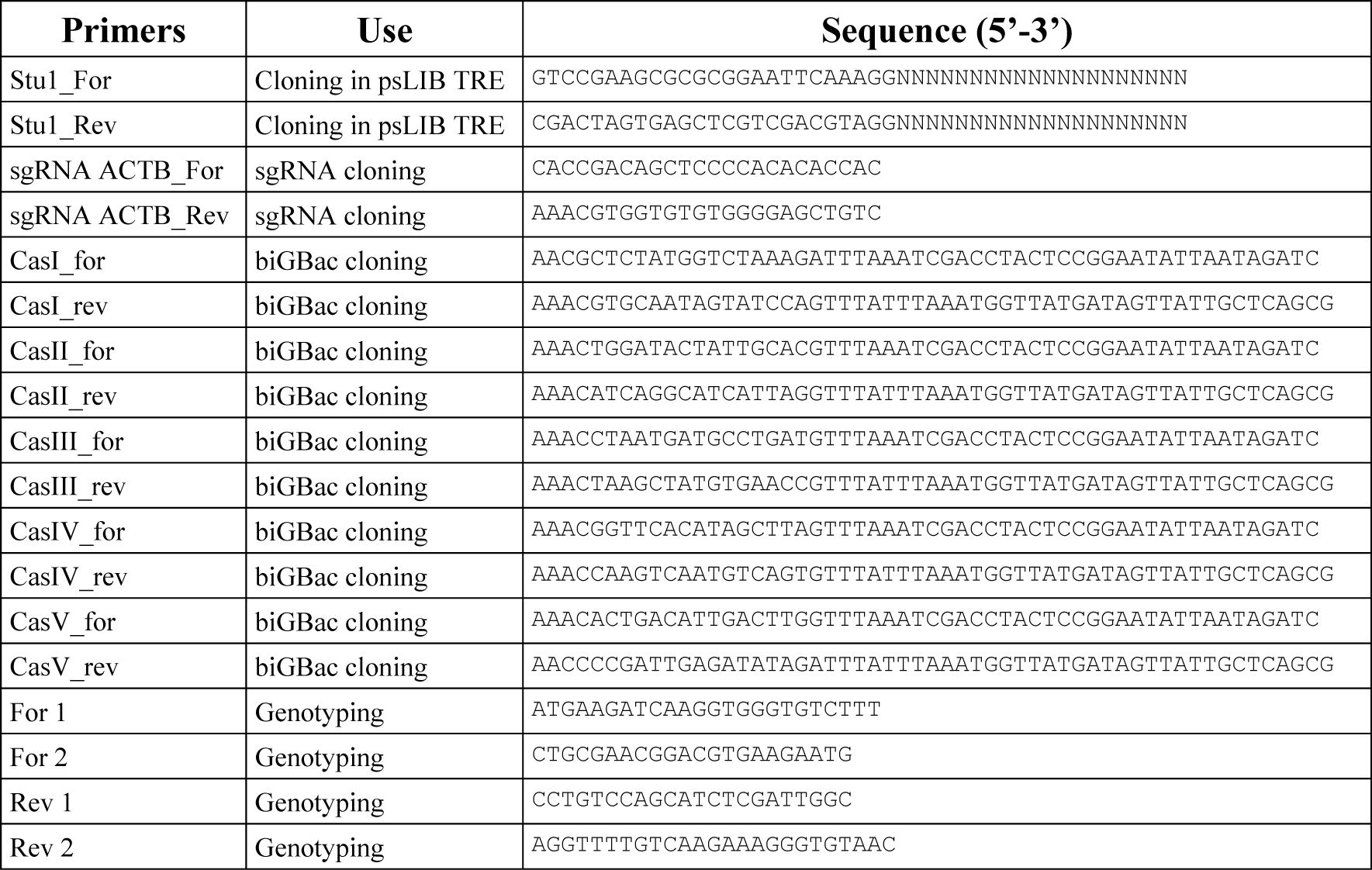
biGMamAct cloning and genotyping primers.

### Equipment and software

- Eppendorf tubes (Dutscher, cat. no. 033511)
- PCR tubes 200 µL (VWR, cat. no. 732-0680)
- Centrifuge (Eppendorf, model no.5424R; rotor no. FA-45-24-11)
- Centrifuge (Eppendorf, model no.5804R; rotor no. A4-44)
- Nanodrop 2000 (Thermo Fisher, model no. ND-2000)
- Dry heat block (Stuart, model no. SBH130D)
- Vortex mixer (Carl Roth, model no. P505.1)
- Falcon tubes (15 and 50 mL; Dutscher cat. nos,352096 and 352098)
- Mini-Sub Cell GT Horizontal Electrophoresis System, with mini-gel caster (Bio-Rad, cat. no. 1704467)
- Mr. Frosty™ (Thermo Fisher, cat. no. 5100-0001)
- PowerPac Basic Power Supply (Bio-Rad, cat. no. 1640303)
- ProFlex PCR System (ThermoFisher, cat. no. 4484073)
- GelDoc XR+ system with Image Lab Software (BioRad, cat. no. 1708195) with XcitaBlue conversion screen (BioRad, cat. no. 1708182)
- Incubator for bacterial growth (Memmert, model IN30)
- Shaker for bacterial growth (Infors HT, model Ecotron)
- HERAcell™ C0_2_ incubator (ThermoFisher, cat. no. 50116048
- Bead bath (Lab Armor, cat.no. A1254302)
- Class II Biological Safety Cabinet MSC-Advantage (ThermoFisher, cat. no.51025411)
- Glass slide (Epredia™ AA00000102E01MNZ20)
- Coverslip Ø 20 mm (Dutscher, cat. no. 140539)
- Inverted transmitted light microscope (Carl Zeiss, model Axio Vert.A1)
- Confocal microscope (Leica TCS SP5 AOBS with laser emissions at 405nm, 458nm, 476nm, 488nm, 496nm, 514nm, 561nm, 594nm and 633nm)
- SnapGene (Version 7.2)
- LAS software (Leica)
- FIJI software (https://imagej.net/software/fiji/)

### Procedure

The following biGMamAct procedure describes how to assemble several genetic modules, namely a construct for conditional and simultaneous tagging of 5 cellular compartments and the generation and isolation of clonal population of a cell line harboring an 18.3-kb-long DNA cargo stably integrated through CRISPR/Cas9.

### **A** Molecular biology. Shuttle and Donor plasmids assembly

The following steps describe how to clone genes of interest into the shuttle vector psLIB and how to assemble several of them to generate single pDONOR vectors suitable for Cas9 mediated HITI-based insertion at the *ACTB* locus.

#### *In silico* design of modules cloning into biGMamAct shuttle vectors and pDONOR assembly ● TIMING 1-2h

We recommend designing all cloning steps computationally Design sLIB TRE backbone

1 Select biGMamAct shuttle vectors considering their different promoters (CMV, EF1α, TRE, or U6)
2 Design cloning oligos matching your cloning method of choice (restriction enzyme or Gibson/LIC). The same multi cloning site (MCS) was inserted in every biGMamAct shuttle plasmids to ease the cloning or exchange of gene of interest (GOI). **▲CRITICAL** If considering involving 2^nd^ round of assembly using biGMamAct, which relies on Pme1 and involves psBIG2 series, validate absence of Pme1 site in the sequences of interest and design silent mutation leading to its removal if any. Of note: we very rarely find this site as hindering cloning despite the huge number of constructs handled at our facility.
3 Validate every cloning steps *in silico* using a dedicated software (SnapGene, ApE^52^…)
4 Order necessary oligos and/or restriction enzyme as required by *in silico* design

#### Cloning GOIs into psLIB TRE shuttle plasmid ● TIMING 3 d

5 psLIB TRE backbone digestion by StuI In a 0.2 mL test tube mix: 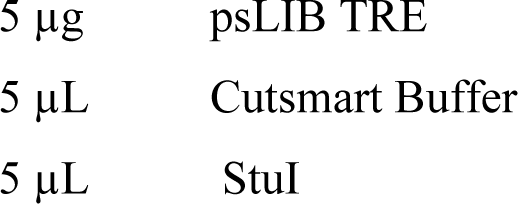 H_2_0 MQ to 50 µL The digestion was incubated at 37 °C for 2 hours.
6 PCR amplification of GOIs. For each PCR amplification mix the following in a 0.2mL test tube: 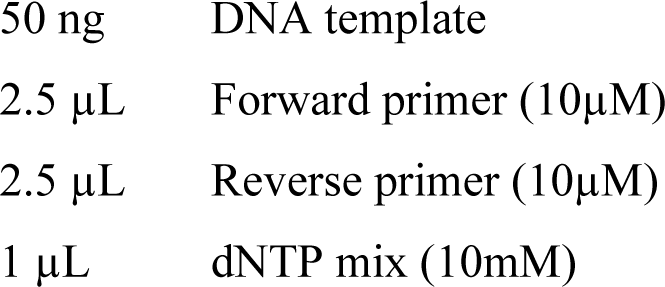 10 µL Q5 polymerase buffer 5X 10 µL Q5 High GC enhancer 5X 0.5 µL Hot Start Q5 High-Fidelity DNA Polymerase H_2_O MQ to 50 µL Run the PCR reactions in a thermocycler with the following parameters: 98 °C 30 seconds 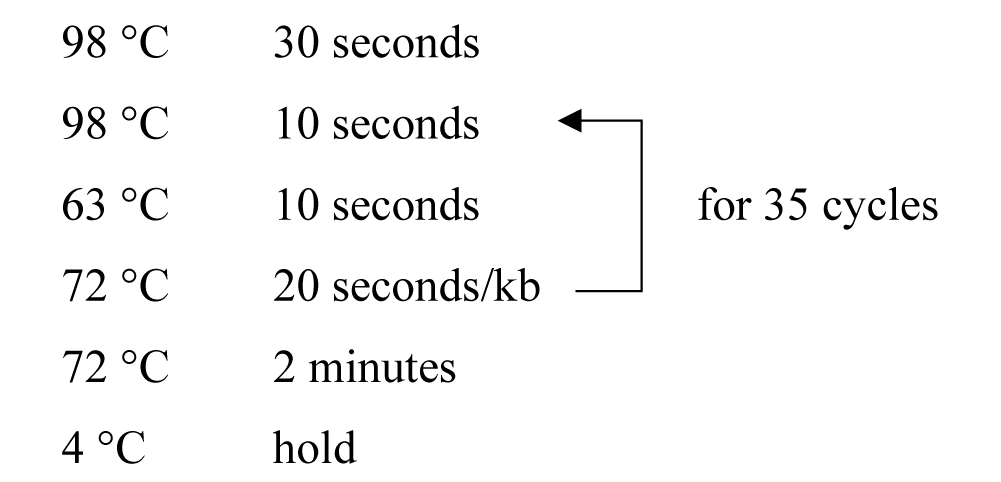 GOIs, i.e., mito-mCherry, eYFP-Tubulin, H2B-IRFP720, GTS-mTagBFP and mTFP1-Actin were respectively amplified by PCR from Addgene 206256, 206254, 206253, 206259 and 206255. **▴CRITICAL** For PCR amplification we use the Q5® High-Fidelity DNA Polymerase (NEB) but other can be picked such as Phusion. **▲CRITICAL** To enable LIC, design forward primer including the 25 bases upstream of the StuI site in psLIB TRE and the reverse primer including the complementary sequence of the 25 bases located downstream of the StuI site (**Table 2**). ° **PAUSE POINT** PCR reactions can be stored at −20 °C up to several months
7 DNA fragment isolation Orange gel loading dye 6X (NEB) was added to the StuI digestion and DpnI treated PCR reactions which were subsequently loaded onto an agarose gel (1% in TAE) supplemented with GelRed (1:10000). 1 kb Plus DNA ladder (NEB) was use as ruler and the gel was run for 30 minutes at 130 V. The bands at 3751 bp (linearized psLIB TRE), 845 bp (mito mCherry), 2169 bp (eYFP tubulin), 1394 bp (H2B iRFP720), 1017 bp (GTS mTagBFP) and 1907 bp (mTFP actin) were visualized using a blue light transilluminator, cut out from the gel with blade and tweezer and purified using QIAquick gel extraction kit (Qiagen). ° **PAUSE POINT** PCR fragments can be stored at −20 °C up to several months
8 Gibson assembly. Genes of interest were then cloned using Gibson cloning into the StuI linearized psLIB TRE vector. For each assembly, mix into a 0.2 mL test tube on ice: 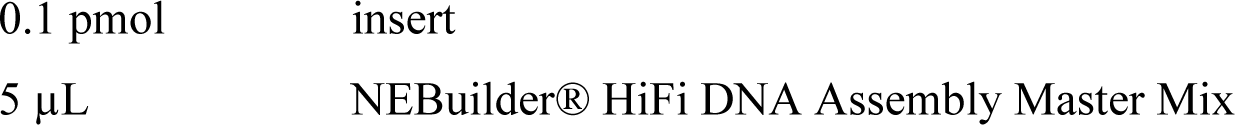 H_2_O MQ to 10 µL Incubate reactions for 15 minutes at 50 °C in a thermocycler. ° **PAUSE POINT** Assembly reactions can be stored at −20 °C up to several months
9 Transform the 10 µL reaction into Top10 chemically competent cells and leave the recovery happen in LB for 1 hour at 37 °C. Plate the transformed cells onto LB-Agar + gentamycin and incubate overnight at 37 °C.
10 The day after pick 3 clones for each construct to isolate the DNA (QIAprep spin Miniprep kit). Measure isolated DNA concentration using a Nanodrop. Screen the DNA assembly through unambiguous restriction digest followed by Agarose gel analysis and send a couple for validation by sequencing.

#### Molecular assembly of pDONOR 6colors ● TIMING 3 d

11 psBIG1a *ACTB*T2ATetP2AP-mAGH linearization using SwaI In a 0.2 mL test tube mix: 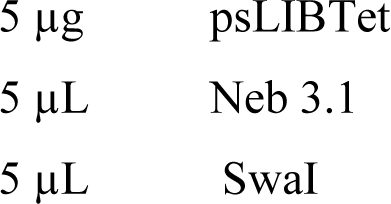 H_2_O MQ to 50 µL The digestion was incubated at 25 °C for 2 hours.
12 PCR amplification of cassettes of interest cloned in psLIB Tet. Our psLIB TRE shuttle plasmid derives from the pLIB of the biGBac system, ie.: individual cassette contained inside can be amplified using standard primers^21^ (**Table 2**) for efficient and reliable multi fragment assembly into our recipient psBIG1a *ACTB*T2ATetP2AP-mAGH harboring a SwaI site flanked with the biGBac α and ω sequences (**Fig 2**). Each individual cassette cloned in psLIB TRE were PCR amplified as described in step 6. Briefly cassette encoding for mito-mCherry was amplified using Cas1-For / Cas1-Rev primers, eYFP-Tubulin using Cas2-For / Cas2-Rev, H2B-iRFP720 using Cas3-For / Cas3-Rev, GTS-TagBFP using Cas4-For / Cas4-Rev and mTFP1-actin using Cas5-For / Cas5-Rev.
13 Add directly into each PCR reaction 0.5 µL DpnI for template digestion to exclude false positive during the next cloning steps. Incubate DpnI treatments for 15 minutes at 37 °C. **▴CRITICAL** it is highly recommended treating with Dpn1 any PCR reaction done using biGMamAct shuttle plasmids prior to their assembly to generate pDONOR ° **PAUSE POINT** PCR reactions can be stored at −20 °C up to several months
14 DNA fragment isolation. Orange gel loading dye was added to the SwaI digestion and DpnI treated PCR reactions which were subsequently loaded onto an agarose gel (1% in TAE) supplemented with GelRed (1:10000). 1 kb Plus DNA ladder was use as ruler and the gel was run for 30 minutes at 130 V. The bands at 6304 bp (linearized psBIG1a *ACTB*T2ATetP2AP-mAGH), 1745 bp (TRE mito mCherry), 3065 bp (TRE eYFP tubulin), 2291 bp (TRE H2B iRFP720), 1913 bp (TRE GTS mTagBFP) and 2800 bp (TRE mTFP actin) were visualized using a blue light transilluminator, cut out from the gel and purified using QIAquick gel extraction kit (Qiagen). **? TROUBLESHOOTING** ° **PAUSE POINT** PCR fragments can be stored at −20 °C up to several months
15 biGBac assembly Each PCR amplified cassettes of interest were then cloned using Gibson cloning into the SwaI linearized psBIG1a *ACTB*T2ATetP2AP-mAGH recipient vector (**Fig 2A**). To do so, mix into a 0.2 mL test tube on ice: To calculate convert 0.06 pmoles of into µL of plasmids of interest, use the following formula:

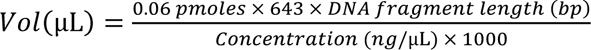 0.06 pmol SwaI linearized psBIG1a *ACTB*T2ATetP2AP-mAGH 0.06 pmol TRE mito mCherry 0.06 pmol TRE eYFP tubulin 0.06 pmol TRE H2B iRFP720 0.06 pmol TRE GTS mTagBFP 0.06 pmol TRE mTFP actin 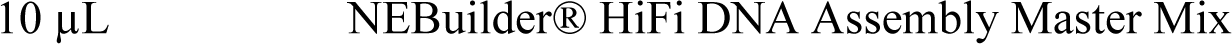 H_2_O MQ to 10 µL Incubate reactions for 1 hour at 50 °C in a thermocycler. **? TROUBLESHOOTING** ° **PAUSE POINT** Assembly reactions can be stored at −20 °C up to several months
16 Transform the 10 µL reaction into Top10 chemically competent cells and leave the recovery happen in LB for 1 hour at 30°C. Plate the transformed cells onto LB-Agar + gentamycin and incubate at 30°C overnight.
17 The day after, pick clones and grow them at 30°C to isolate the DNA (QIAprep spin Miniprep kit). Measure isolated DNA concentration using a Nanodrop. Screen the DNA assembly through unambiguous restriction digest (here SwaI was used for excision of each individual cassette) and send positive for validation by sequencing (**Fig 2B**).

#### Assembly of pX458 *ACTB* ● TIMING 3 d

pX458 (Addgene 48138) is suitable for both expression of SpCas9 and sgRNA. sgRNA cloning is achieved by annealing 2 ssDNA primers and subsequent Golden Gate cloning.

18 BbsI digestion of pX458 In a 0.2 mL test tube mix: 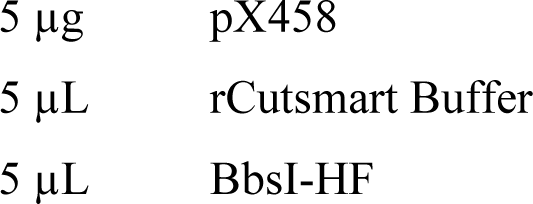 H_2_O MQ to 50 µL Incubate the digestion at 37 °C for 2 hours.
19 Orange gel loading dye was added to the BbsI digestion which was subsequently loaded onto an agarose gel (0.8% in TAE) supplemented with GelRed (1:10000). 1 kb Plus DNA ladder was used as ruler and the gel was run for 60 minutes at 130 V. The bands at 9280 bp (digested pX458) was visualized using a blue light transilluminator, cut out from the gel and purified using QIAquick gel extraction kit (Qiagen). ° **PAUSE POINT** DNA fragments can be stored at −20 °C up to several months. **? TROUBLESHOOTING**
20 Annealing of ssDNA primers Prepare the annealed oligo master mix by pipetting in a 1.5 ml eppendorf: 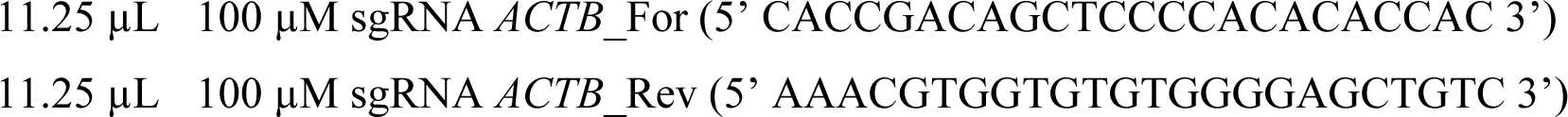 2.5 µL of 10X Annealing Buffer Mix, spin and incubate at 95 °C for 5 minutes. Turn the heat block off and leave it cool naturally with the eppendorf inside. When the temperature reaches 50-40°C, let it cool at room temperature. Alternatively, a gradient can be set up in the PCR machine for gradual cooling. ° **PAUSE POINT** This mix can be used directly or stored at −20 °C up to several months.
21 Annealed oligos dilution The Annealed oligos master mix must be diluted 400 times in 0.25 X Annealing Buffer: in a new 1.5 mL eppendorf mix: 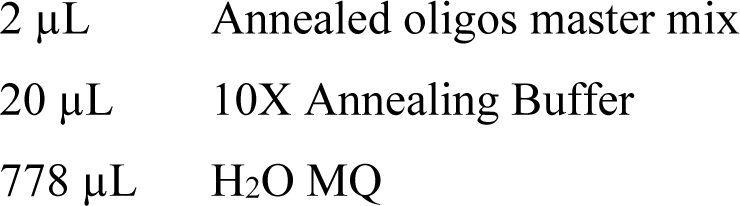
22 Cloning of pX458 *ACTB* Ligate annealed oligos into BbsI cut pX458 plasmid in a 1.5 mL Eppendorf tube: 200 ng BbsI digested pX458 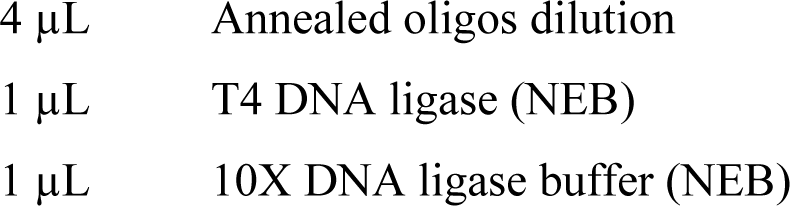 H_2_O MQ to 10 µL Incubate 1 hour at 25 °C or at RT. ° **PAUSE POINT** Ligation reaction can be stored at −20 °C up to several months.
23 Transform 5 µL of the ligation mix into Top10 chemically competent cells and leave the recovery happen in LB for 1 hour at 37 °C.
24 Plate the transformed cells onto LB-Agar + ampicillin (100 µg/mL) and incubate at 37 °C overnight.
25 Pick 4 clones and grow them overnight at 37 °C for subsequent isolation of the DNA (QIAprep spin Miniprep kit). Measure isolated DNA concentration using a Nanodrop. Screen the clones through unambiguous restriction digest (eg. BbsI / EcoRV or equivalent) and send a couple of positive for validation by sequencing. **? TROUBLESHOOTING** **B Cell culture**
26 Cell maintenance: Culture HEK293T cells (ATCC CRL 3216) in DMEM high glucose + 10% FCS + pen/strep (complete medium) in a 37 °C, 5% CO_2_ incubator and passage them every 3-4 days by trypsinization when reaching 90% confluency. **▲CRITICAL** it is highly recommended starting stable cell line establishment using low passage cell (<10), tested against mycoplasma. **▲CRITICAL** For each of the following trypsinization steps, confirm complete cell dissociation under a light microscope.

#### Transfection ● TIMING 30 min

27 Seed 1.5 million HEK293T cells into one well of a 6-well plate in 1.5 mL complete medium.
28 Pipette in a 1.5 mL Eppendorf tube 50 µL DMEM (without FBS and pen/sterp) 800 ng pDONOR 6 colors 250 ng pX458 *ACTB* Mix by quick vortexing.
29 In a separate 1.5 mL Eppendorf tube prepare 50 µL DMEM (without FBS and pen/strep) 3 µL LipoD293 reagent Mix by quick vortexing.
30 Transfer the 53 µL DMEM + LipoD293 solution into the 1.5 mL Eppendorf containing DMEM+DNA and mix by pipetting. **▲CRITICAL** For proper DNA/LipoD293 complexes formation, pipette DMEM + LipoD293 into DMEM + DNA and not the opposite.
31 Incubate 10 minutes at RT to allow DNA/LipoD293 complex formation. Add drop by drop the whole content of the tube onto the cells and incubate the plate in a 37 °C, 5% CO_2_ incubator.
32 The day after replace the medium with 2 mL complete medium.

#### Establishment of the stable cell line ● TIMING 15 d

33 Puromycin and hygromycin selection. 72 hours post transfection dissociate the cells, pellet and seed them all into a 10 cm dish with 12 mL complete medium containing 0.5 µg/mL puromycin and 100 µg/mL hygromycin. Every 2-3 days, monitor cell selection and exchange the medium keeping 0.5 µg/mL puromycin and 100 µg/mL hygromycin. After applying selection for 15 days, HEK293T cells should be recovering. **▲CRITICAL** From now on, keep maintaining edited HEK293T in complete medium containing 0.5 µg/mL puromycin and 100 µg/mL hygromycin. **▴CRITICAL** If biGMamAct is applied onto other cell types than HEK293T it is mandatory to perform dose/response analysis for puromycin and hygromycin prior applying selection for stable cell line generation **? TROUBLESHOOTING**

#### Recovered polyclonal cells and clonal seeding ● TIMING 10 d

34 When reaching ∼30% confluency in a 10 cm dish, dissociate and count the cells (approx. 5 million cells).
35 Maintain 0.5-1 million cells in culture in a 10 cm dish in complete medium containing 0.5 µg/mL puromycin and 100 µg/mL hygromycin. Such culture may be used for putative clonal seeding if required by any reasons.
36 Pellet 3 million cells and resuspend them into 2 mL complete medium + 10% sterile DMSO. Distribute 2 x 1 mL into 2 cryovials and freeze the cells using a Mr. Frosty™.
37 Save a 0.5-1 million cell pellet and place it at −20 °C for further genomic DNA extraction and genotyping. This is the “pool” sample. Of note: if targeting genomic locus other than *ACTB*, one can start optimizing genotyping PCR conditions with the “pool” sample and non-edited WT cells (step 57).
38 Seed 2 plates for clonal isolation containing 12mL complete medium containing 0.5 µg/mL puromycin and 100 µg/mL hygromycin with 300 cells in each.
39 After 5 days replace medium by fresh medium containing 0.5 µg/mL puromycin and 100 µg/mL hygromycin. Growing clones should be visible using a light microscope. Leave the plate for 5 more days until individual clones are visible by eye.

#### Clone picking ● TIMING 2 h

(The following steps describe the picking of 48 clones)

40 Add 20 µL Trypsin in 48 wells of a 96-well plate.
41 Remove medium from the dish containing individual clones and slowly add 8 mL PBS against the dish wall (not directly onto the cells).
42 Set a p10 micropipette to 8 µL. With a sterile tip detach, by gently scrubbing, one single clone while slowly aspirating it – once fully detached suck it up.
43 Transfer the aspirated clone into one well of the 96-well plate containing trypsin and resuspend up and down.
44 Repeat this for 47 clones. **▲CRITICAL** Replace PBS every 10 pickings or from the moment clones/cells are floating. **▲CRITICAL** From now on, when working with individual clone, change tip (or aspirating pipette) to avoid cross contamination.
45 Once done, incubate the 96-well plate in a 37 °C, 5% CO_2_ incubator for 2 minutes.
46 Meanwhile, add 400 µL complete medium containing 0.5 µg/mL puromycin and 100 µg/mL hygromycin into each well of a 48-well plate.
47 Add 100 µL complete medium containing 0.5 µg/mL puromycin and 100 µg/mL hygromycin into each well of the 96-well plate. Resuspend few times the picked clones changing tip.
48 Transfer the 128 µL from each well of the 96-well plate into wells of the 48-well plate refilled with 400 µL complete medium.
49 Replace medium after 24 hours with 400 µL complete medium containing 0.5 µg/mL puromycin and 100 µg/mL hygromycin.

#### Clonal expansion and genotyping ● TIMING 8 d

Here, 24 clones will be expanded and genotyped to highlight the power of the method but users can work with less.

50 After 3-5 days, identify the grown/recovered clones under a light microscope (i.e. approx. confluent) and dissociate them. Briefly, remove medium, gently wash the cells with PBS prior adding 50 µL trypsin per well. After 5 minutes in a 37 °C, 5% CO_2_ incubator add 950 µL complete medium containing 0.5 µg/mL puromycin and 100 µg/mL hygromycin per well and resuspend cells by pipetting up and down 10 times prior transferring them all (1 mL) into 12-well plates for amplification.
51 After 3-4 days, i.e. when almost confluent, dissociate the clones using 100 µL trypsin per well and 1 mL complete medium containing 0.5 µg/mL puromycin and 100 µg/mL hygromycin to quench the reaction.
52 Resuspend cells by pipetting up and down 10 times and transfer for each clone, 1 mL of resuspended cell into a sterile 1.5 mL Eppendorf.
53 Add back 1 mL complete medium containing 0.5 µg/mL puromycin and 100 µg/mL hygromycin into each well still containing 100 µL of cells and place plates in a 37 °C, 5% CO_2_ incubator. Once positive clones identified through genotyping, expand, freeze and characterize them from these plates.
54 Pellet clonal cell populations, here 24 tubes, 3 minutes at 300 *g*. Discard the medium and save the cell pellets for genomic DNA (gDNA) extraction and subsequent genotyping. ° **PAUSE POINT** Cell pellets can be stored a −20 °C up to several weeks.
55 gDNA extraction of individual clones and the “pool” (step 34) can be achieved through different methods. Here, we use the Monarch® Genomic DNA Purification Kit (New England Biolabs) and follow the manufacturer recommendation. Final gDNA elution is done in 30 µL elution buffer and yields typical gDNA concentration between 50-100 ng/µL.
56 Genotyping of 5’-, 3’- and WT junctions is achieved by running PCRs on extracted gDNA of each individual clone. Couple of primers used for genotyping the 5’-, the 3’- and the WT junctions are For1 /Rev1, For2/Rev2 and For1/Rev2 respectively (**Table 2**). **▴CRITICAL** If genotyping other edited junctions (i.e. targeting different genomic locus), it is highly recommended optimizing PCR conditions by working only with “pool” sample and non-edited WT cells (step 38).
57 For every clone, “pool” and HEK293T WT set three PCRs for mapping each junction by mixing in a 0.2 mL test tube: 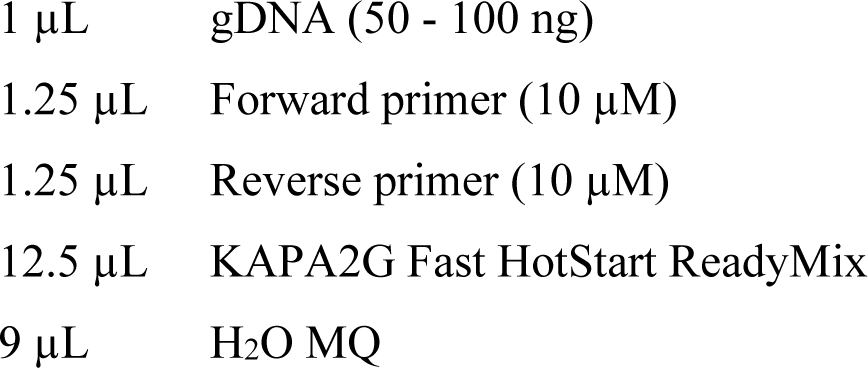 **? TROUBLESHOOTING**
58 Run the PCR in a thermocycler: 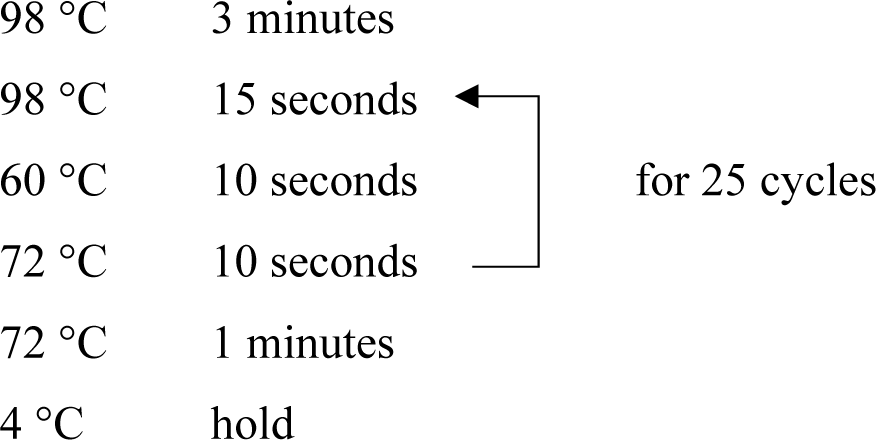 **? TROUBLESHOOTING**
59 Load PCR reactions onto an agarose gel (1% in TAE) supplemented with GelRed (1:10000). Run the gel for 30 minutes at 130 V and use 1 kb Plus DNA Ladder as ruler.
60 Identify correct integration through visualization of bands at 526 bp and 386 bp for the 5’ and 3’ junction respectively. Heterozygotes still show a band at 313 bp outlining the persistence of the WT junction (**Fig 3**).

#### Functional Validation ● TIMING 3 d

The following section is not directly part of our method and should be adapted dependent on the genes of interests and on the intended application. Here, we present a proof of concept illustrating the strength of biGMamAct: six genetically encoded modules (including one located into the psBIG1a backbone) can easily be assembled in one single step for subsequent precise and highly efficient integration at the *ACTB* locus while still retaining their functions.

61 Doxycycline induction of fluorescent reporters. To validate functional DNA integration, positive clones were cultured in complete medium containing 0.5 µg/mL puromycin and 100 µg/mL hygromycin, onto collagen-coated Ø 20 mm glass coverslip. Once reaching ∼50% confluency, medium was replaced by complete medium containing 0.5 µg/mL puromycin, 100 µg/mL hygromycin and 1 µg/mL doxycycline. 24 hours after, medium was removed and cells were washed with PBS prior fixation.
62 Slide preparation. Doxycycline induced cells were fixed using PFA (4% in PBS) for 15 minutes at 37 °C. After PFA removal and two PBS washes, coverslips were mounted onto glass slide using ProLong™ Diamond Antifade Mountant with DAPI. Mounted glass slides were kept in the dark at 4 °C for 24 hours.
63 Imaging. Cells were imaged using a Leica TCS SP5 AOBS confocal microscope. Characteristic fluorescence and localization corresponding to every module inserted into the pDONOR 6colors were monitored and homogenously observed in every cell: mTagBFP at the Golgi apparatus, mTFP-actin (cytoskeleton), mAzamiGreen at the nucleus, eYFP tubulin (microtubules), mCherry at the mitochondria and iRFP720-Histone2B at nucleus (**Fig 4**).

**Figure 4:**
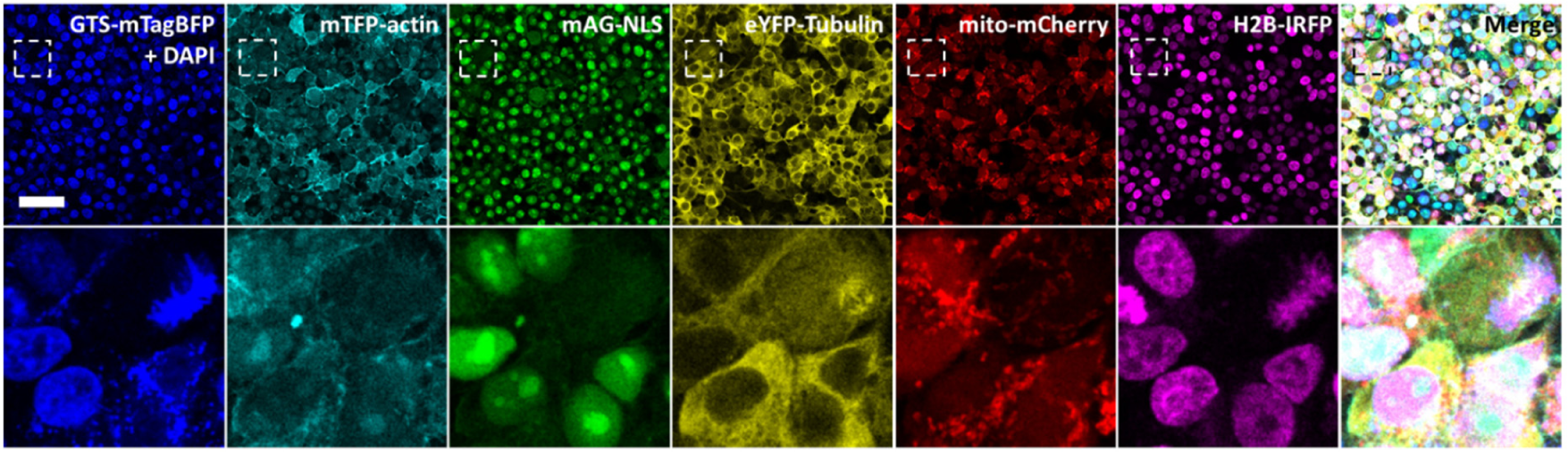
Confocal imaging of HEK293T edited clone. Pictures evidencing in all cells stable integration of large and functional DNA payloads through homogeneous tunable expression and correct subcellular localization of GTS-mTagBFP (Golgi), mTFP1-Actin (cytoskeleton/adhesion), mAG-NLS (nucleus), YFP-Tubulin (microtubules), mito-mCherry (mitochondria) and H2B-iRFP (histones) on top of DAPI staining. Lower panel is a zoom into larger field of view from above (dashed box) and showing different appearance of tagged cellular compartments during cell cycle. For example, during metaphase the golgi apparatus is fragmented and no more visible, actin filaments are mostly visible at the cell periphery, GFP-NLS is evenly distributed within le cell, microtubules form spindle spanning from centrosomes to chromosomes, mitochondria are highly fragmented and H2B colocalizes with the condensed chromosomes gathered at the metaphase plate. Scalebar, 50 μm.

## Anticipated results

Using biGMamAct, we have demonstrated that 6 coloring modules of different subcellular compartments with tunable expression can be assembled in one single step despite their highly repetitive sequences. After pDONOR 6colors assembly, we have underlined the strength of biGMamAct through CRISPR/Cas mediated insertion at the *ACTB* locus with high efficiency and precision. Relying on the sgRNA/spCas9 and HITI tandem for stable integration of >18kb DNA cargo at *ACTB* locus showed very high efficiency (100% homozygotes). Our biGMamAct system, with its portfolio of shuttle plasmids, allowed for the first time the precise knock-in of 18.3 kb functional DNA payload. Indeed, we speculate involvement of Doxycycline-inducible promoters most probably helped lifting silencing barriers especially true for large DNA integration. We anticipate that the use of biGMamAct will be a promising tool for characterization of unelucidated complex biological processes notably through *in situ* structural investigations^2,35^, and will unlock hitherto insuperable barriers in synthetic biology for metabolic engineering in mammalian cells^19,32,33^, as recently done in prokaryotes^53^ or plants^54^,.

### TROUBLESHOOTING

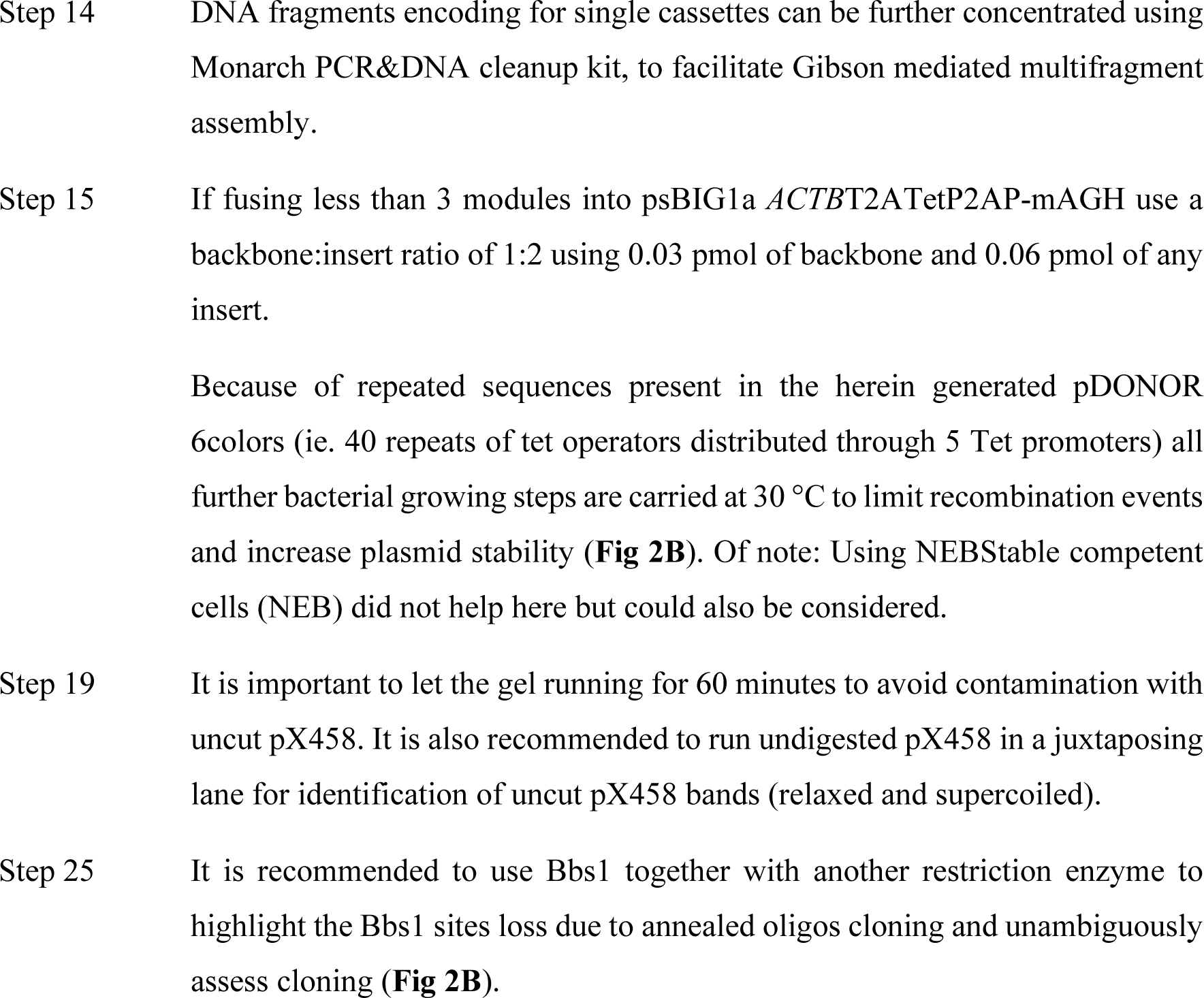
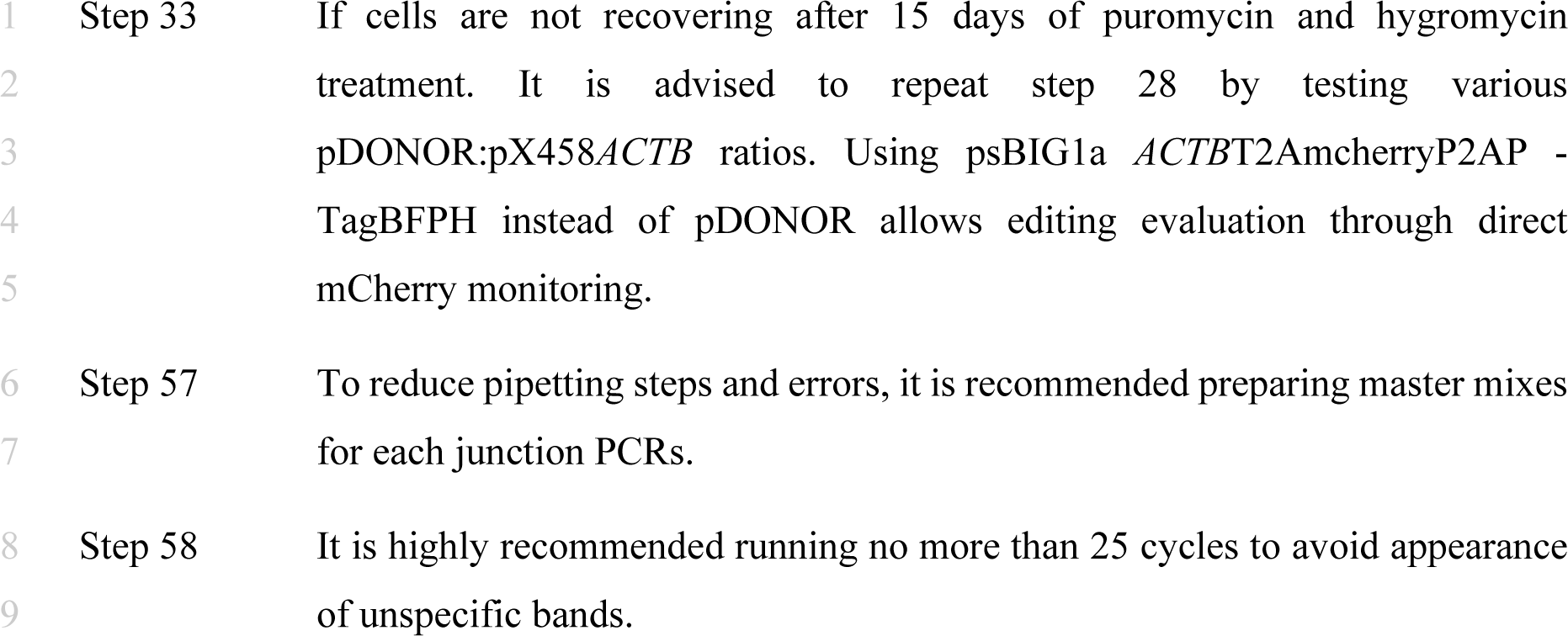

### TIMING

The timing notes listed in the Procedures refer to the timing typically required by a trained researcher to implement the protocol (PhD student or more senior researcher). The timing is also critically dependent on the exact cell line of interest used and integration efficiency may depend on its state of ploidy. We indicate the timing suitable to perform the experiments using the same reagents and cell line that we have used, as listed in the Materials and Reagents section.

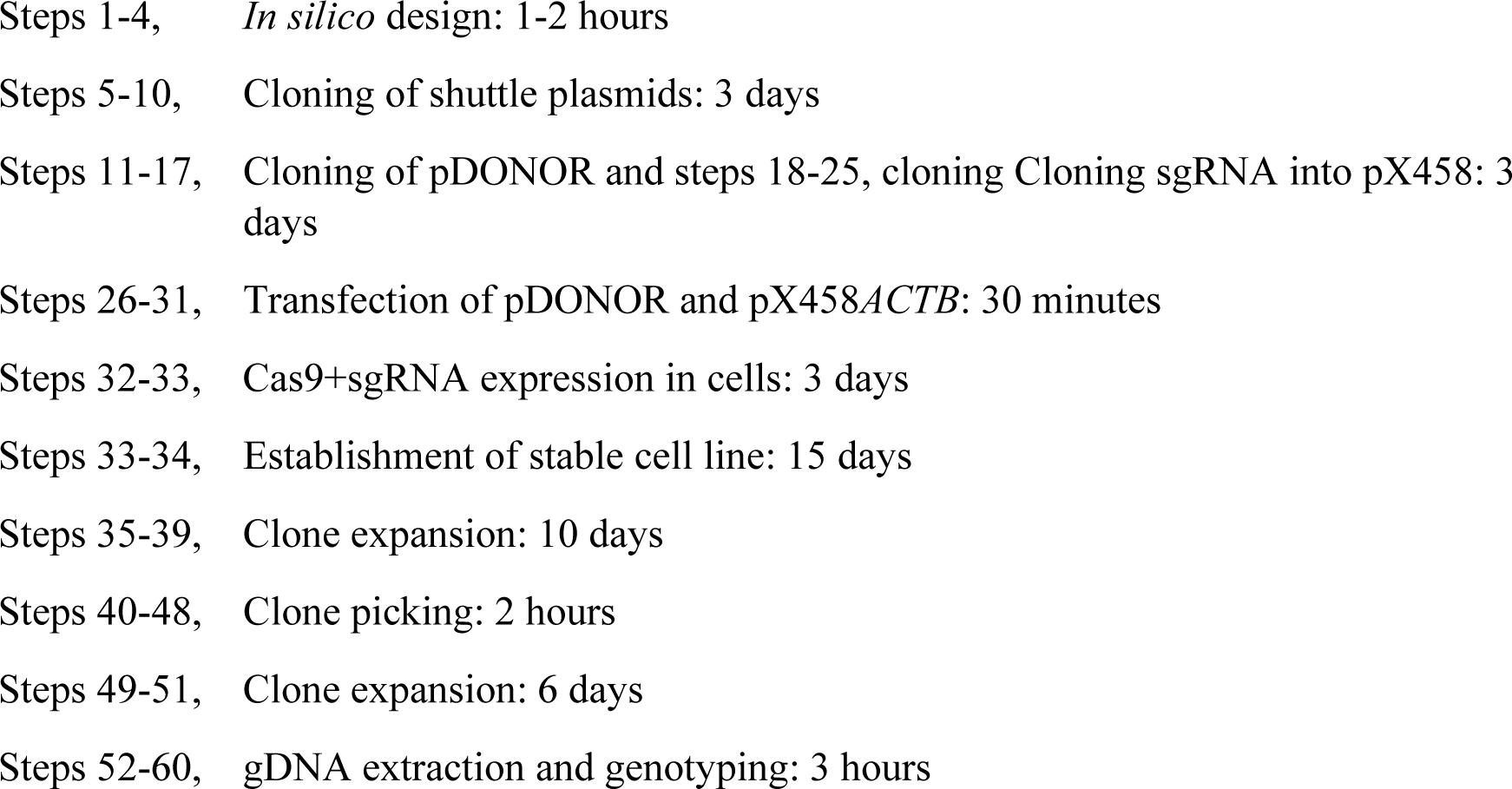

## Author contributions statements

MP and MM conceived the project. MM supervised the research. MP designed and performed all experiments. MP wrote the first draft of the manuscript. MP and MM wrote and approved the final draft of the manuscript.

## Acknowledgments

We thank Prof. Kristina Djinovic-Carugo for critical reading of the manuscript and all members of EMBL Grenoble for helpful discussion.

## Data availability statement

All vectors used in this protocol will be available from the corresponding authors on request.

